# RNA-guided gene drives can efficiently and reversibly bias inheritance in wild yeast

**DOI:** 10.1101/013896

**Authors:** James E. DiCarlo, Alejandro Chavez, Sven L. Dietz, Kevin M. Esvelt, George M. Church

## Abstract

Inheritance-biasing “gene drives” may be capable of spreading genomic alterations made in laboratory organisms through wild populations. We previously considered the potential for RNA-guided gene drives based on the versatile CRISPR/Cas9 genome editing system to serve as a general method of altering populations^1^. Here we report molecularly contained gene drive constructs in the yeast *Saccharomyces cerevisiae* that are typically copied at rates above 99% when mated to wild yeast. We successfully targeted both non-essential and essential genes, showed that the inheritance of an unrelated “cargo” gene could be biased by an adjacent drive, and constructed a drive capable of overwriting and reversing changes made by a previous drive. Our results demonstrate that RNA-guided gene drives are capable of efficiently biasing inheritance when mated to wild-type organisms over successive generations.

## Introduction

Gene drives have the potential to address diverse ecological problems by altering the traits of wild populations. As ‘selfish’ genetic elements, they spread not by improving the reproductive fitness of the organism, but by increasing the odds that they themselves will be inherited. Because this inheritance advantage can counteract the fitness costs associated with the drive itself or with adjacent genes carried along with it, they are theoretically capable of ‘driving’ unrelated traits through populations over many generations^1^.

Inheritance-biasing is a common strategy in nature^2^. One elegant class of inheritance-biasing genes spreads by cutting homologous chromosomes that do not contain them, thereby inducing the cellular repair process to copy them onto the damaged chromosome by homologous recombination (Fig. 1A). This process is known as ‘homing’^3^. The best-known homing endonuclease gene is *I-SceI*, whose product cuts the gene encoding the large rRNA subunit of *S. cerevisiae* mitochondria. Most are capable of homing with extremely high efficiencies; I-SceI is correctly copied 99% of the time^4^.

**Figure 1.**
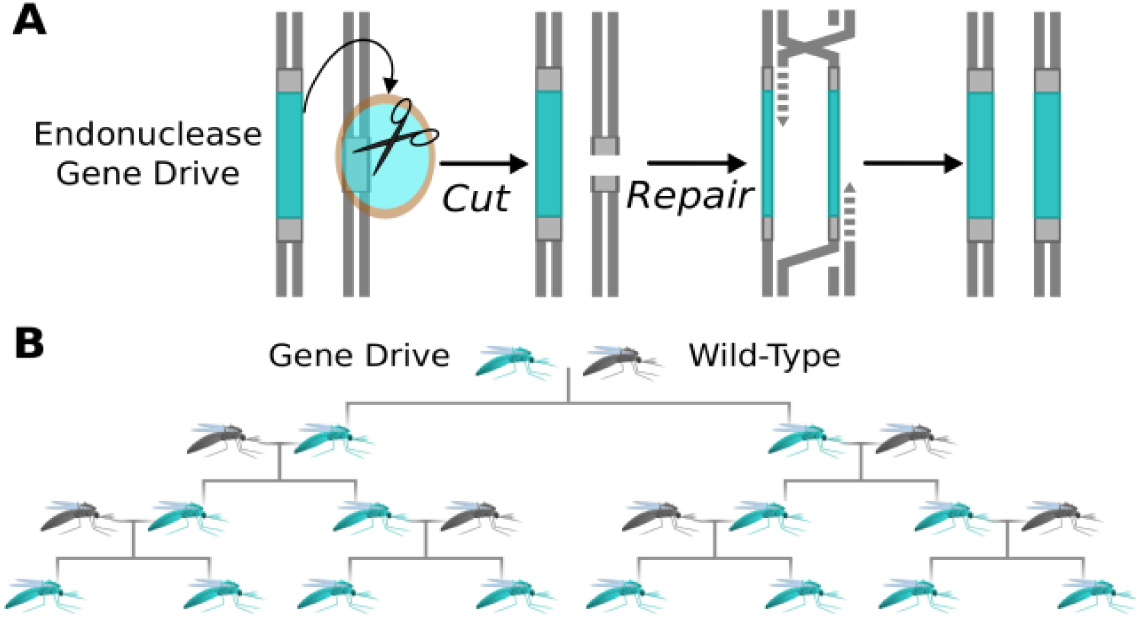
Mechanism and population-level effect of endonuclease gene drives. (A) Homing endonucleases cut competing alleles, inducing the cell to repair the damage by copying the endonuclease gene. (B) By converting heterozygous germline cells into homozygotes containing two copies (teal), gene drives increase the odds that they will be inherited and consequently spread themselves and associated changes through wild populations (grey).

Austin Burt first suggested that homing endonucleases might be used to construct synthetic gene drives capable of altering wild populations of multicellular organisms in 2003 (Fig. 1B)^5^. Laboratories subsequently reported that the *I-SceI* endonuclease gene exhibited homing in transgenic laboratory populations of mosquitoes^6^ or fruit flies^7,8^ with an I-SceI recognition site inserted into the corresponding locus. However, gene drives based on homing endonucleases are constrained by the difficulty of retargeting these enzymes to cleave useful sequences within wild-type genomes^9^.

The recent development of the CRISPR nuclease Cas9, which cleaves target sequences specified by “guide RNA” molecules, has enabled scientists to edit the genomes of diverse species^10–20.^ Because Cas9 can apparently be used to edit any gene, the question of whether it can be used to gene drives capable of spreading those changes through wild populations is highly relevant. We previously detailed the theoretical potential for RNA-guided gene drives to alter wild populations, including an evolutionary analysis, novel drive architectures, and containment measures robust to human error^1^. However, these new architectures and control strategies have not yet been validated. Indeed, whether Cas9 can bias inheritance at all remains unknown, raising the question of whether initiating public discussions and engaging in regulatory reform^21^ are immediately necessary. We sought to address these issues by constructing several types of RNA-guided gene drives and quantifying their ability to bias inheritance in the yeast *S. cerevisiae*.

## Results

Because gene drives have the capacity to alter native populations, we took stringent precautions to prevent the accidental escape of our gene drives into wild yeast. We first employed a method of molecular containment^1^ in which we split our Cas9 based gene drive system into two physically separate genetic parts: an episomally encoded Cas9 gene and a drive element encoding the guide RNA. This allowed us to avoid creating a self-sufficient inheritance-biasing cassette while still targeting wild-type yeast strains. Though simple, this form of molecular containment is not vulnerable to human error; even if drive-containing yeast were to escape into the wild, the required Cas9 episomal plasmid would rapidly be segregated away from the drive element, rendering the drive inoperative.

To directly measure the efficiency of Cas9 gene drives in yeast, we used the *ADE2* gene as a visual marker^22^. If red *ade2Δ* haploids are mated with cream-colored wild-type haploids, the resulting heterozygous diploids inherit one functional copy of *ADE2* and are cream-colored. When these diploids undergo meiosis and reproduce via sporulation, half the resulting haploids inherit the broken copy and are consequently red; the other half inherit the intact copy and are cream-colored (Fig. 2A).

**Figure 2.**
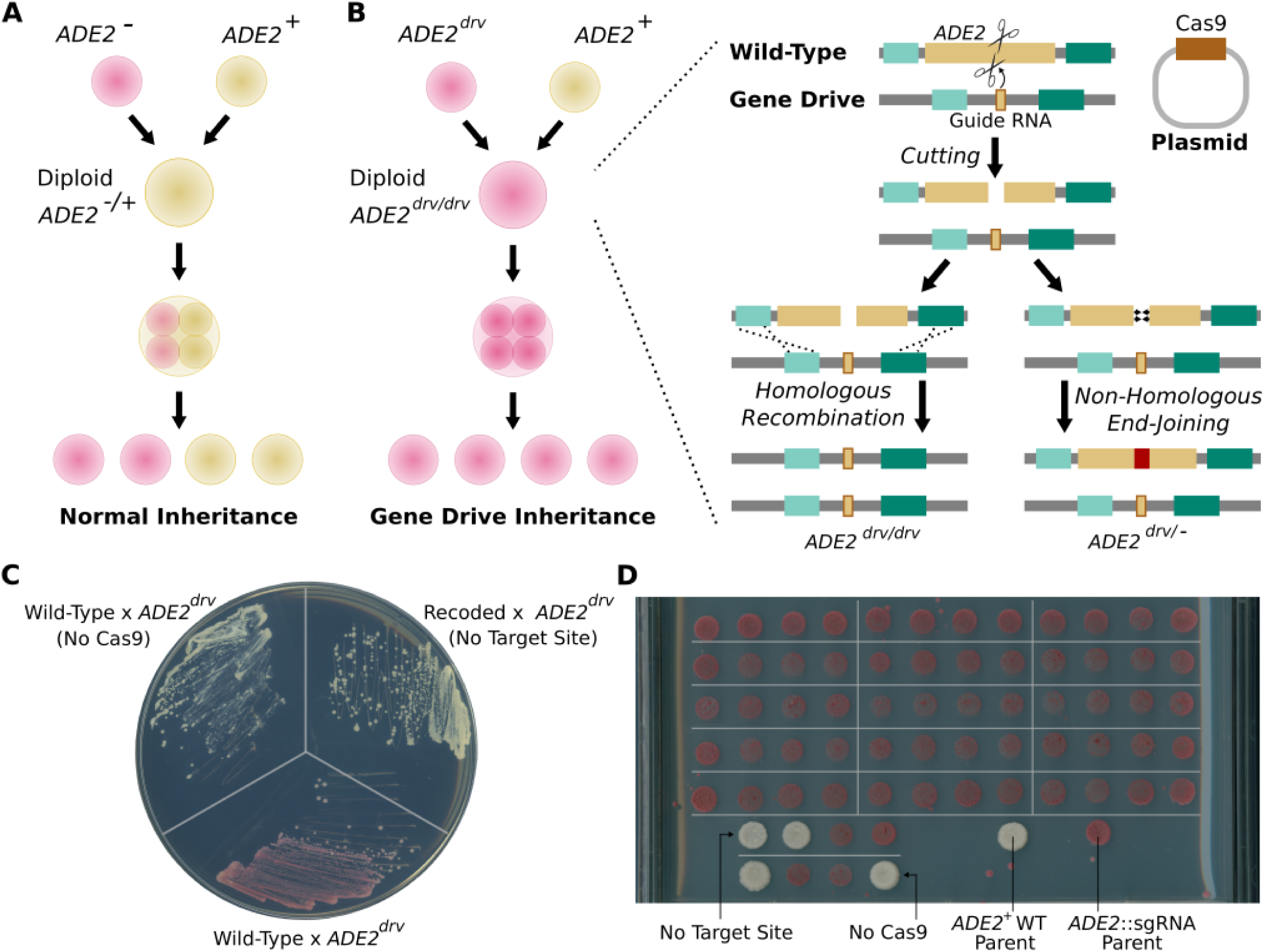
Biased inheritance of *ADE2* is readily visible in *S. cerevisiae*. (A) Mutations in *ADE2* generate a red phenotype on adenine-limiting media due to the buildup of red pigments. Mating a red mutant haploid to a wild-type haploid produces cream-colored diploids, which yield 50% red and 50% cream-colored progeny upon sporulation. (B) When haploids with a gene drive targeting *ADE2* mate with wild-type haploids in the presence of Cas9, cutting and subsequent replacement or disruption of *ADE2* produces red diploids that upon meiosis yield exclusively red progeny. (C) Diploids produced by mating wild-type and *ade2*::sgRNA gene drive haploids yield cream-colored colonies in the absence of Cas9 or when the target site is removed by recoding but uniformly red colonies when both are present, demonstrating Cas9-dependent disruption of the wild-type *ADE2* copy. (D) Spores from 15 dissected tetrads produce uniformly red colonies on adenine-limited plates, confirming disruption of the *ADE2* gene inherited from the wild-type parent. In the absence of the target site or Cas9, normal 2:2 segregation is observed.

But if the red haploids encode a functional gene drive in place of *ADE2*, it will cut and replace the intact *ADE2* locus inherited from the wild-type parent, yielding red diploids. Following meiosis, all haploid progeny will inherit one of the two gene drive alleles and will also be red (Fig. 2B). Thus, the cutting efficiency of a gene drive that replaces *ADE2* can be assessed by mating drive-containing haploids to wild-type haploids, selecting for diploids and counting the fraction that are red.

We built a Cas9-based gene drive targeting *ADE2* by placing a guide RNA against the wild-type *ADE2* gene in place of the endogenous *ADE2* locus. We mated these red *ade2*::sgRNA haploids to wild-type yeast of the opposite mating type in the presence or absence of the Cas9 plasmid and plated on media that selects for diploids. Nearly all diploid colonies were red when the Cas9 plasmid was present, indicating highly efficient cutting of the *ADE2* copy inherited from the wild-type parent (Fig. 2C). As expected, we did not observe red diploid colonies in the absence of Cas9, demonstrating that the drive only functions in populations encoding both Cas9 and guide RNA.

To verify that the *ADE2* alleles from drive-containing diploids were indeed lost, mated diploids were sporulated and their resultant haploid progeny were examined. Upon dissecting 18 *cas9*+ diploids, we observed a perfect 4:0 ratio of red:cream haploids, confirming that all copies of the *ADE2* locus were disrupted. In contrast, 18 cream-colored *cas9*-diploids yielded a 2:2 red: cream ratio, indicating normal inheritance of the inactivated drive and the wild-type *ADE2* allele (Fig. 2D).

To determine whether *ADE2* disruptions in red diploids were the result of successful copying of the drive element, we sequenced the 72 haploids derived from dissected *cas9*+ diploids. All sequenced colonies contained intact drives without additional mutations, indicating that drive mobilization was efficient and occurred at high fidelity. At the time of submission and preprint release, this was the first example of a synthetic endonuclease gene drive that biases its own inheritance when mated to a wild-type organism.

We next tested whether RNA-guided gene drives could be designed to bias the inheritance of not only the minimal drive element, but also any closely associated “cargo” gene whose spread through an existing population may be desirable. As a proof of principle, we inserted the *URA3* gene *in cis* to the *ade2*::sgRNA drive element. *URA3* allows laboratory modified yeast strains to grow in the absence of uracil supplementation (Fig. 3A). We mated these *URA3*-containing drive haploids to wild-type haploids in the presence of an episomal Cas9 plasmid, selected diploids (all of which were red), sporulated them, and dissected 18 tetrads. As was the case for the original *ADE2* gene drive, all of the sporulated haploid cells formed red colonies. Crucially, all grew normally when replica plated on uracil deficient media, indicating that *URA3* was efficiently copied with the drive (Fig. 3B).

**Figure 3.**
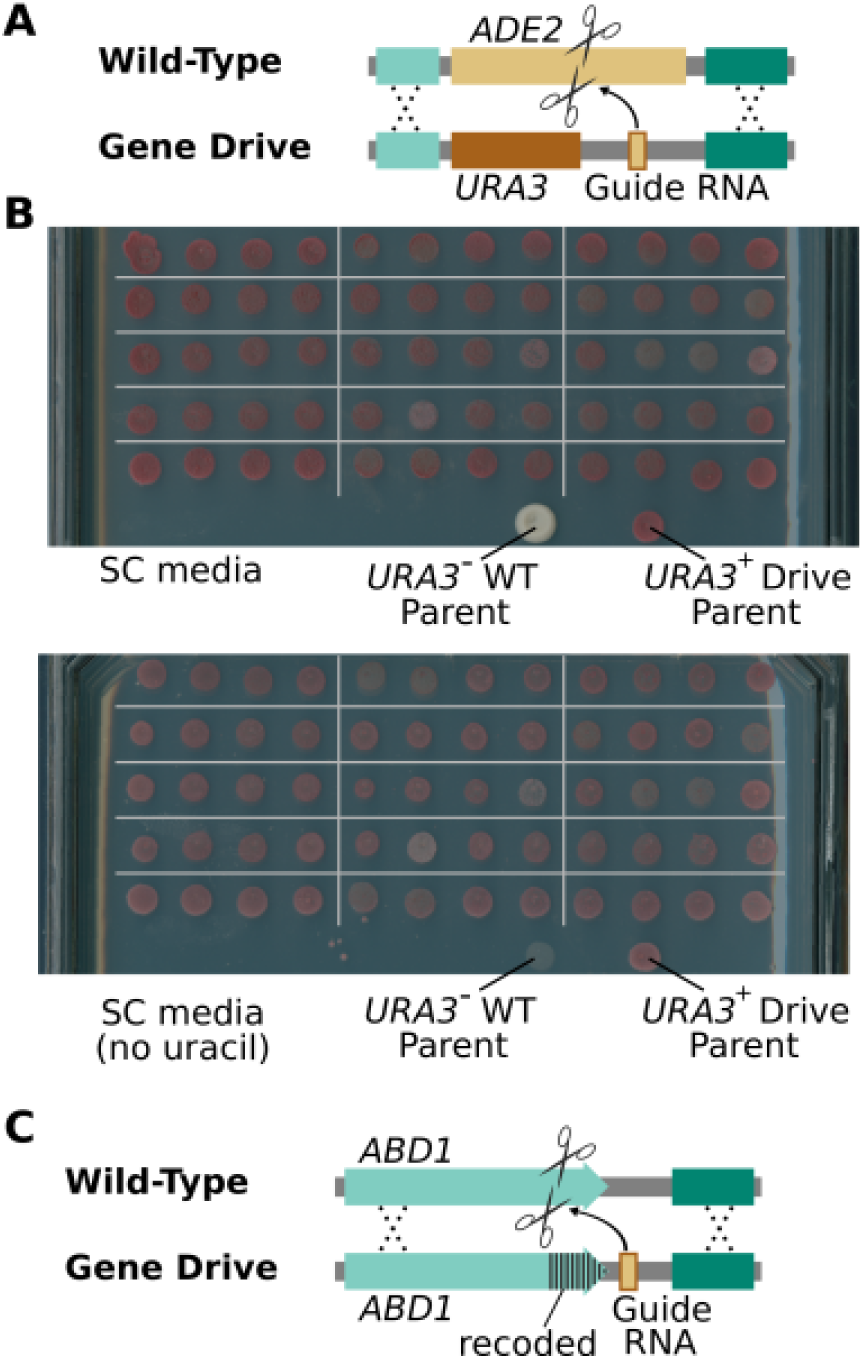
Gene drives and cargo genes remain intact upon copying and can spread by targeting both non-essential and essential genes. (A) The *ADE2*-targeting gene drive was modified to carry *URA3* as a cargo gene. (B) Diploids produced by mating wild-type *URA3-*haploid yeast with haploids encoding the gene drive carrying *URA*3 were sporulated and tetrads dissected to isolate colonies arising from individual spores. Pictures are spores from 15 of these tetrads. All grew when replica-plated onto plates lacking uracil, demonstrating that the drive successfully copied *URA3* in all diploids. (C) Depiction of a gene drive designed to cut and recode the 3′ end of the essential *ABD1* gene.

We subsequently sought to determine whether gene drives can not simply disrupt a gene, but recode it and leave the function intact. A gene drive targeting an essential gene in this manner should be more evolutionarily stable than one that cuts a non-essential gene such as *ADE2*, which can be readily blocked by mutations that mutate or remove the target site^1^. We consequently built a drive targeting the essential *ABD1* gene (Fig. 3C)^24^. We mated this haploid strain, which has a recoded *ABD1* allele upstream of the guide RNA, to wild-type cells in the presence of Cas9. We then selected diploids, sporulated the cells, dissected 18 of them, and sequenced the 72 resulting segregants. All contained the drive element and recoded *ABD1* locus, thereby validating our proposed essential gene recoding architecture.

All of our earlier experiments involved matings between haploids of the same strain. We were curious whether our gene drives could be copied into a diverse group of wild *S. cerevisiae* strains at equal efficiency. These strains may vary in some of the many factors determining gene drive copying efficiency, such the types of repair machinery available to the cell at the time of the cut, the chromatin state of the locus, and the degree of homology flanking the double-strand break generated by the drive. We correspondingly mated *ADE2* drive-containing haploids with 6 phylogenetically and phenotypically diverse wild-type strains of haploid *S. cerevisiae*^25^.

To more accurately measure the efficiency with which all of these gene drives were copied in various backgrounds, we performed quantitative PCR on populations of the resulting diploids using one set of primers specific to the drive and another set designed to amplify either wild-type alleles or those disrupted by non-homologous end-joining.

The mean fraction of diploid chromosomes containing the *ADE2* gene drive was over 99% regardless of wild-type parent strain (Fig. 4), attesting to the robustness of the drive in diverse backgrounds. Of note, addition of the *URA3* cargo gene did not appreciably change this efficiency. The drive that targets and recodes the essential *ABD1* gene was similarly efficacious.

**Figure 4.**
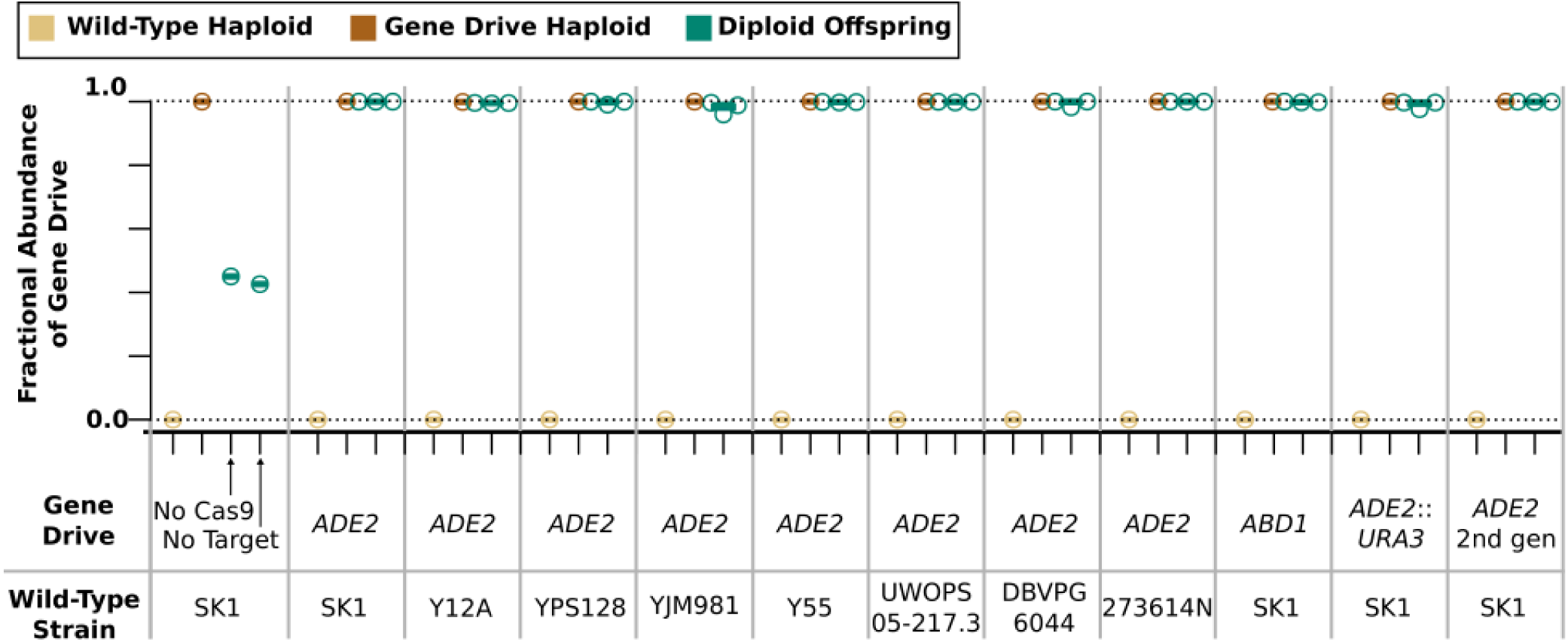
The extent of inheritance-biasing induced by various gene drives in diverse yeast strains as measured by quantitative PCR. Results depict the relative abundance of wild-type and drive-containing alleles in diploids arising from matings between SK1 haploids bearing gene drives and diverse wild-type haploid strains. “No Cas9” and “No Target” refer to haploid cells containing the *ADE2* drive mated to wild-type haploids in the absence of Cas9 or to an otherwise wild-type strain with Cas9 that has a mutation in the targeted sequence that blocks cutting. “2nd gen” refers to the haploid progeny of an earlier mating.

We next determined whether RNA-guided gene drives could bias inheritance over successive generations. Because *S. cerevisiae* reproduces mostly through asexual division and only a subset of the population sporulate in the laboratory even when induced to undergo meiosis^23^, experiments to determine the long-term population-level efficiency of gene drives are difficult to perform and interpret. As a surrogate for population takeover experiments, we sought to measure the performance of successive copying events by a single drive element. Towards this goal, we mated the haploid progeny of the *ADE2* gene drive to wild-type haploids, selected for diploids, and ran quantitative PCRs. The constructs biased inheritance at the same efficiency in the second generation as they did in the first (Fig. 4, far right).

We designed the previously described gene drives to be incapable of autonomous spread due to their requirement for exogenously supplied Cas9 (Fig. 2C–D, Fig. 4). To determine whether an autonomous drive encoding the large *cas9* gene is similarly efficient, we constructed one such drive targeting a recoded synthetic sequence within *ADE2* (Fig. 5A). We further sought to determine whether the loss of *ADE2* function induced by this drive might be undone using a “reversal drive”^1^. We consequently built a second autonomous gene drive to cleave the first autonomous *ADE2*-disrupting drive and subsequently restore an intact copy of the *ADE2* gene (Fig. 5B). Quantitative PCR demonstrated that both of these autonomous drives were copied at an average efficiency over 99% (Fig. 5D). Together, these results demonstrate the efficacy and reversibility of autonomous Cas9-based gene drives.

**Figure 5.**
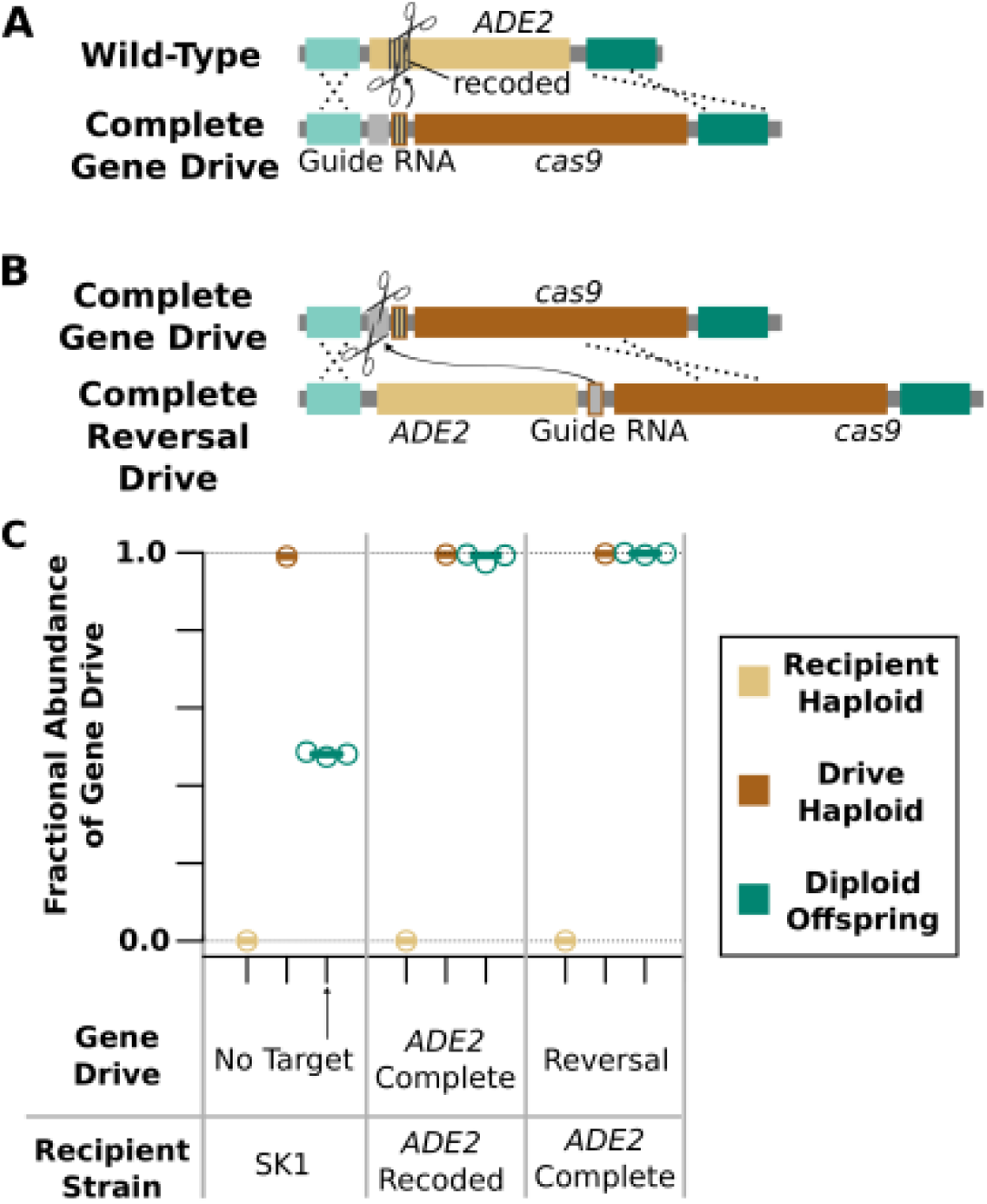
Essential, autonomous, and reversal gene drives. (A) A complete and autonomous gene drive that cuts and replaces the recoded *ADE2* gene. (B) A reversal drive that cuts the autonomous drive and restores *ADE2*. (C) Quantitative PCR results depicting the relative abundance of wild-type and drive-containing alleles in diploids arising from matings between SK1 haploids bearing the above gene drives and wild-type SK1 yeast. <u>Fig. 8-S1</u> depicts sporulated haploid progeny of the reversal drive crossed with the autonomous *ADE2*-disrupting gene drive.

## Discussion

Our discovery that Cas9 can bias inheritance in diverse wild yeast strains over successive generations at very high efficiency demonstrates that RNA-guided gene drives can function in eukaryotic organisms as predicted. By itself this does not guarantee the success of gene drives in other organisms as the rate of homologous recombination varies between species and is known to be particularly high in yeast. However, the fact that we observed inheritance biasing rates equal to or exceeding that of the natural *I-SceI* homing endonuclease gene is highly promising. Because a drive based on I-SceI efficiently initiated copying following successful genome cutting events in transgenic *Anopheles gambiae* mosquitoes that contained an I-SceI recognition site^6^ and Cas9 typically cuts more efficiently than does I-SceI^7,25^, our findings suggest that RNA-guided gene drives will be highly effective at suppressing ^5,26,27^ or spreading antimalarial alleles^28,29^ through populations of this important malaria vector. Similarly, RNA-guided gene drives are likely to be more effective than those based on I-SceI, zinc-finger nucleases, or TALENs^7,8,30^ due to their ability to efficiently cut multiple sequences without requiring highly repetitive elements.

The success of our *ABD1* gene drive demonstrates the feasibility of targeting and recoding genes important for fitness, a strategy expected to improve the evolutionary stability of gene drives^1^. We recommend that future efforts seeking to build gene drives intended for eventual release adopt this same approach. On a similar note, our successful test of a reversal drive construct strongly suggests that such drives should be constructed in tandem with any gene drive capable of enacting a specific change in a wild population as a safeguard against accidental escape.

More generally, our findings suggest that yeast may prove a useful platform for swiftly testing RNA-guided gene drive architectures before moving them into multicellular organisms. The power of yeast genetics and the ease of genome manipulation will facilitate combinatorial investigations into gene drive optimization. For example, studies might explore how biasing repair pathway choice^31^ affects the efficiency of copying for gene drives of various sizes. Because the factors involved in these pathways are broadly conserved, these experiments could guide gene drive optimization in other organisms^32^.

While highly encouraging for potential gene drive applications, our results also sound a note of caution for subsequent experiments. That our drives were readily copied into a variety of yeast strains collected from all over the world underscores the potential for a single gene drive to affect very large populations. Poor flanking homology is not an effective barrier, at least in *S. cerevisiae*. Moreover, the *ADE2* gene drive took only two weeks to design, build, and test, suggesting that many laboratories are capable of building gene drives in yeast. Since yeast reproduce mainly through asexual division, gene drives would need to be considerably less costly to organismal fitness in order to spread in the wild than would a comparably efficient gene drive in an organism that always reproduces via mating. However, natural endonuclease gene drives such as *I-SceI* do exist within yeast. Whether our gene drives or the typical RNA-guided gene drive will constitute this level of burden is as yet unknown.

It is more difficult to edit the genomes of multicellular model organisms such as *Drosophila* than it is for yeast, and still more difficult to alter those species for which gene drive applications are most likely to be relevant. However, a growing number of laboratories now make heritable alterations in more than a dozen sexually reproducing species. This confluence of factors demands caution. Because synthetic gene drives would alter the global environmental commons, the decision to deploy such a drive *<u>must</u>* be made collectively by society. Any accidental release could severely damage public trust in scientists. As demonstrated by numerous containment breaches involving pathogenic viruses and bacteria, physical methods of containment are always susceptible to human error and should not be exclusively relied upon whenever alternatives are available.

All scientists making heritable alterations with Cas9 should therefore employ non-physical containment methods sufficient to prevent the creation of an RNA-guided gene drive capable of spreading in the wild. Even scientists not intending to work with gene drives should consider taking precautions, since any unintended insertion of the cas9 gene and guide RNAs near a targeted site could generate a gene drive. Fortunately, a simple and costless precaution is both available and already utilized for different reasons by many laboratories: *avoid delivering the* Cas9 *gene on a DNA cassette that also encodes a guide RNA*. As we have shown, guide RNAs alone cannot bias inheritance in the absence of Cas9 and consequently cannot spread through wild populations (Fig. 4B).

We recommend that future gene drive experiments in yeast similarly separate Cas9 from guide RNAs or employ another method of molecular containment, as exemplified by our autonomous gene drives that target synthetic sequences not found in wild genomes. Gene drive experiments in species that always reproduce sexually pose greater risks and consequently should employ additional precautions such using both forms of molecular containment or performing experiments in geographic areas where the organism in question cannot survive – a form of ecological containment^1^. Working in genetic backgrounds that are less likely to escape the laboratory and mate, such as wingless flies in the case of *Drosophila*, may also be prudent.

In conclusion, our demonstration of diverse gene drive architectures enabling Cas9-mediated inheritance biasing in wild *S. cerevisiae* can guide efforts to build RNA-guided gene drives in other organisms and underscores the urgent need for precautionary control strategies, inclusive public engagement, and regulatory reform^21^ in advance of real-world applications.

## Methods

### Plasmids and genomic cassettes

Gene drive cassettes were synthesized from gBlocks (Integrated DNA Technologies, Coralville, IA) and inserted into SK1 cells via Cas9-mediated genome modification as follows. Guide RNAs for each drive were cloned into p416-Cas9 containing plasmids with expression driven by the SNR52 promoter^18^. 60 base pair homology arms to the target locus were added on both ends of the gene drive cassette via PCR and 5 ug of PCR product was co-transformed with the p416-Cas9-gRNA plasmids. Correctly integrated gene drives were verified by sequencing and p416-Cas9-gRNA plasmids were cured using 5-Fluoroorotic Acid (FOA) selection.

To create the *URA3-*containing *ADE2* gene drive, the *ADE2* gene drive was cloned next to the *Candida albicans URA3* gene in the pAG60 plasmid. The entire URA3 cassette and gene drive were PCR amplified and inserted using Cas9-mediated genome modification into the *ADE2* locus of haploid SK1 cells.

The recoded C-terminus of the *ABD1* gene and corresponding gene drive were synthesized as a gBlock to remove homology and generate mutations in the seed sequence via synonymous changes. The TEF1 terminator was inserted at the 3’end of the recoded *ABD1* gene between the gene and the gRNA as *ABD1* shares a terminator with the *VHC1* gene. The entire cassette was integrated into the haploid SK1 genome using Cas9-mediated genome modification.

The ADE2 gene was recoded by cotransforming a double stranded oligonucleotide and a p416 plasmid containing Cas9 and a gRNA targeting the ADE2 region to recode. The oligonucleotide silently recoded the ADE2 gene and included an orthogonal target and PAM sequence. The complete gene drive (Cas9 and gRNA, targeting the recoded ADE2 gene) was generated by cloning a gRNA into the p416-Cas9 plasmid. An orthogonal genomic target was also included in the complete gene drive to later be targeted by the reversal drive. The Cas9 and gRNA linear construct was amplified by PCR using the same homology arms as the sole gRNA gene drive construct. The construct was co-transformed into By4723 cells with the plasmid it was amplified from and the cells were selected for uracil prototrophy. Correct integrations were screened via colony PCR. This plasmid was later removed using FOA.

The reversal drive (Cas9 and gRNA integrated upstream of the ADE2 gene) was generated by cloning an alternately encoded gRNA into a p414 plasmid containing Cas9. This alternatively endoded gRNA contains less homology to previously used gRNAs and would reduce the chance of unwanted recombination when used to replace the complete gene drive. This gRNA targets a 20bp region inserted with the complete gene drive. The TRP1 gene with the Cas9 and gRNA were PCR amplified with homology arms to the 5’ region of the ADE2 and the product was transformed into SK1 A cells. Cells were selected for tryptophan prototrophy and screened via PCR for correct integrations.

The p416-Cas9-gRNA plasmid (conferring uracil prototrophy) is a variant of the previously described p414-Cas9-gRNA plasmid (conferring tryptophan prototrophy)^18^ (Addgene #43802). One or the other was used in each mating experiment. The pRS413 vector was transformed into select cell types to confer histidine prototrophy as a marker to select for diploid cells.

Strain genotypes:

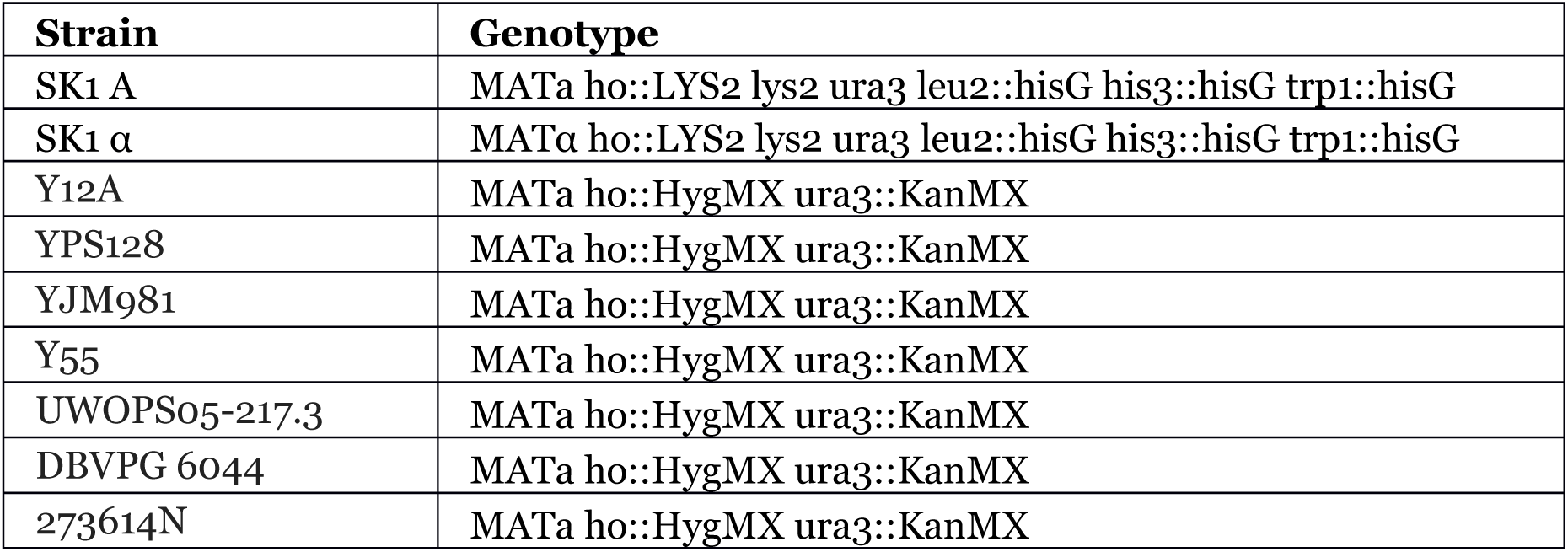

### Ye ast mating experiments

Haploid drive-containing SK1 yeast and haploid wild-type strains of the opposite mating type were mixed in equal amounts in YPAD liquid media and incubated overnight. The resulting diploids were washed in sterile water and plated on selective media for both parental genotypes. The chart below details the specific crosses:

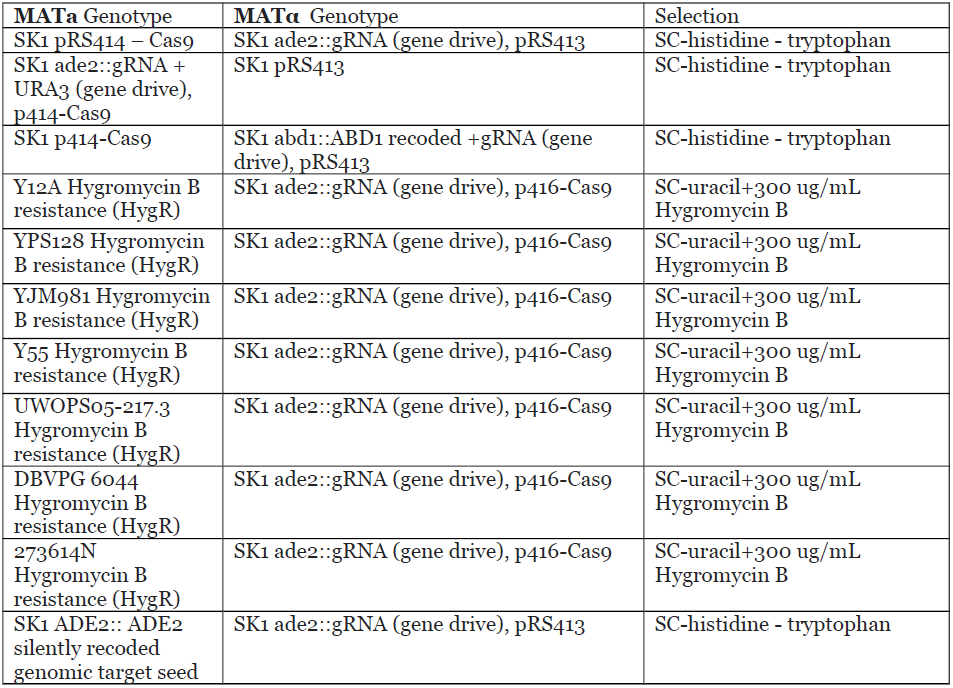
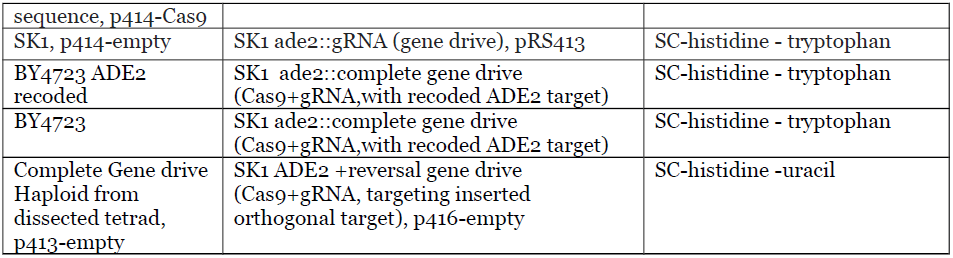

### Sporulation and tetrad dissection

After mating in liquid YPAD and selection for diploids on selection plates, the selection plates were scraped into 10 mL selective media and grown overnight at 30°C. A fresh 5 mL YPAD culture was then inoculated to and OD=0.1 and grown 4-5 hours at 30°C. The entire culture was then washed twice in 10 mL water, inoculated into 2 mL of sporulation media(1% potassium acetate), and incubated at room-temperature for 3 days or until spores were visible. Sporulated cells were suspended in 50 μL of a stock solution of zymolyase (50 μg/mL in 1M sorbitol) and incubated at 30C for 5 minutes, transferred to ice, diluted with 150 μL cold H2O, microdissected using a Zeiss tetrad dissection microscope, and isolated spores grown on YPAD plates.

### Selection for URA3 function

Dissected spores were grown in synthetic complete (SC) media and then spotted onto SC medium as well as SC medium without uracil. To enhance red color, all SC solid media used for plate images contained 0.5 X adenine hemisulfate (final concentration of 0.08 mM).

### Verification of chromosomal segregation

Three genes on chromosome 15 flanking the ADE2 gene were sequenced in two dissected tetrads of the complete gene drive cross. VAM3, TRS33, and DPP1 were amplified using PCR from colonies and sequenced using Sanger sequencing to verify that homologous recombination copied only the gene drive cassette rather than the entire chromosome.

### Quantitative PCR

Candidate primer pairs were designed to amplify short regions specific to each drive or the wild-type sequence replaced by the drive, as well as the *ACT1* gene as a control. All sequences are included in the supplementary information. Genomic DNA was extracted using Method A as described in Looke et al.^33^

KAPA SYBR FAST qPCR Master Mix (2X) was used to perform the qPCR reaction along with 25 ng of genomic DNA. The amplification efficiency and relative specificity of each primer pair were measured by amplifying dilutions of genomic DNA from wild-type and drive haploids, respectively, and the best-performing and well-matched pairs selected for use (see below for all primers used). Quantitative PCR reactions were performed on genomic DNA isolated from each parental haploid as well as from diploids arising from three independent mating events. Three reactions (technical replicates) were performed per sample on a LightCycler 96 machine by Roche.

### Calculations

Results from three technical replicates were averaged for calculations. In order to directly calculate the ratio of alleles before PCR amplification, we first determined the efficiencies of the different primer pairs. Efficiencies were calculated from qPCR runs of serial dilutions (6 orders of magnitude) as:

Efficiency=10^-1/slope^

R^2^ values were higher than 0.99 in all cases except for one pair (ade2::URA3+sgRNA).

The allelic ratios were calculated as:

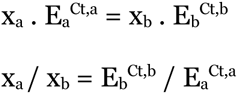

with

x_a_ and x_b_ being the initial concentration of drive and wt DNA,
E_a_ and E_b_ the efficiency of the respective primer pairs and
Ct,a and Ct,b the Ct values for each sample.

Figure 4B was generated using BoxPlotR^34^.

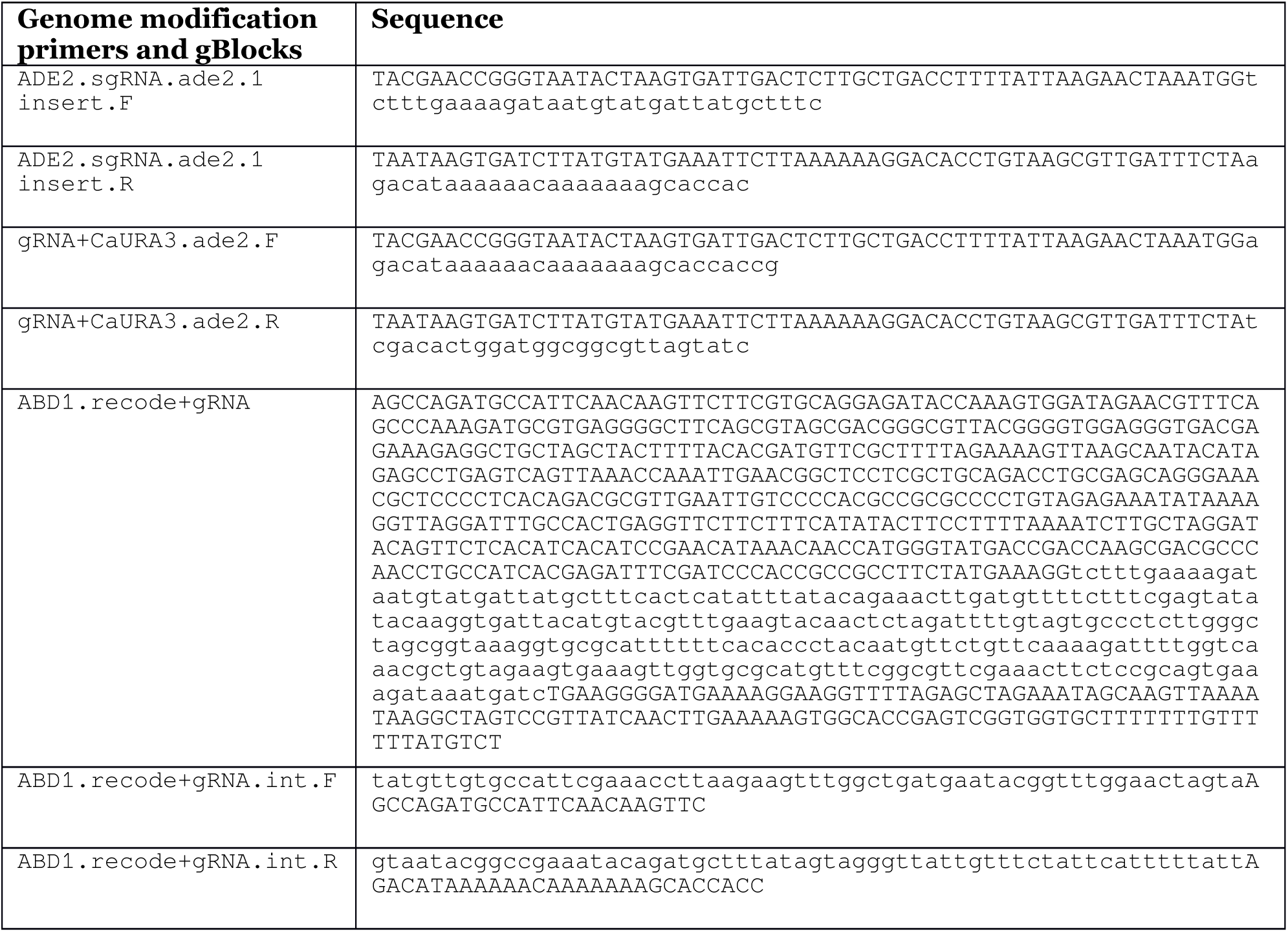

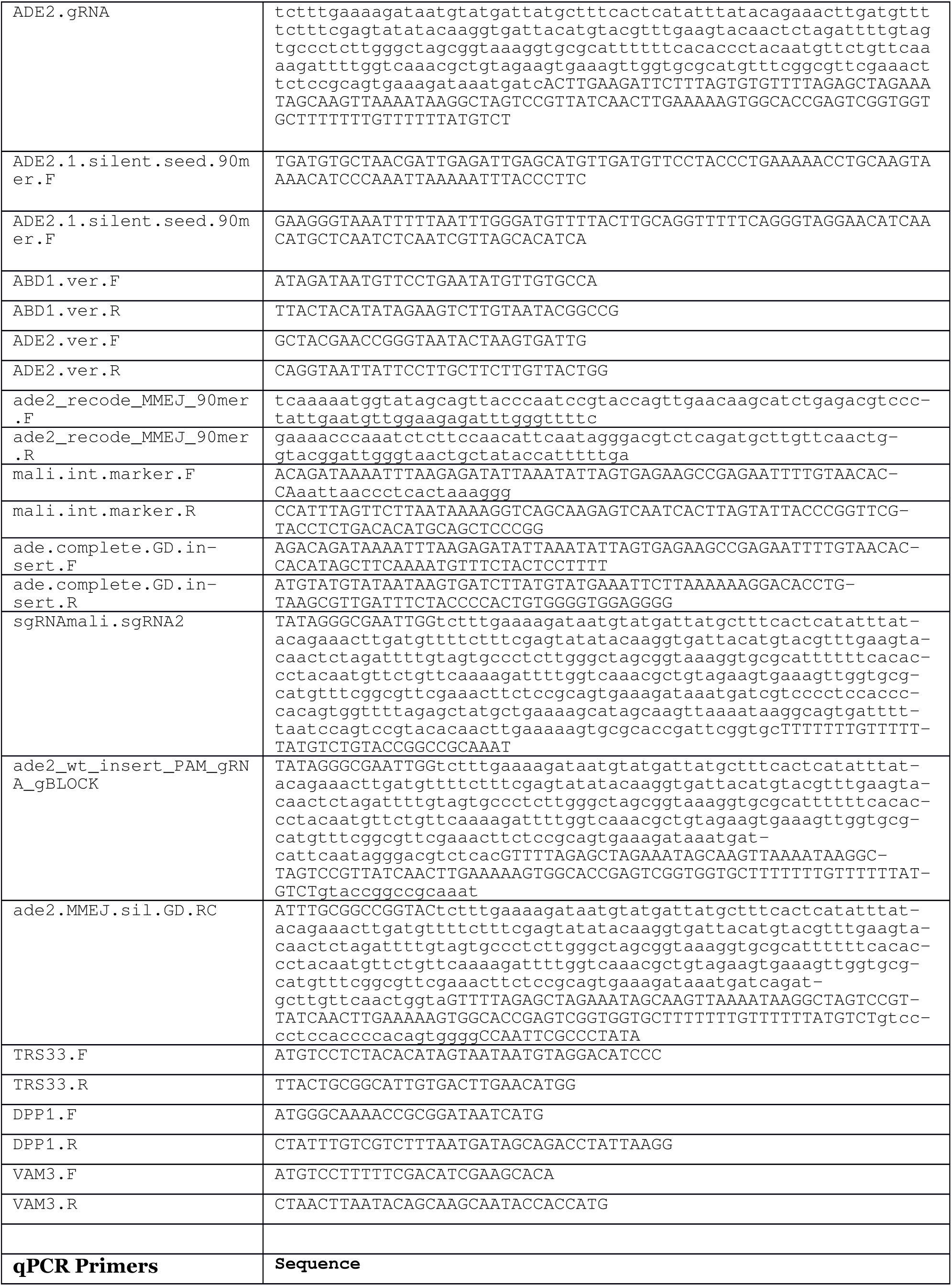

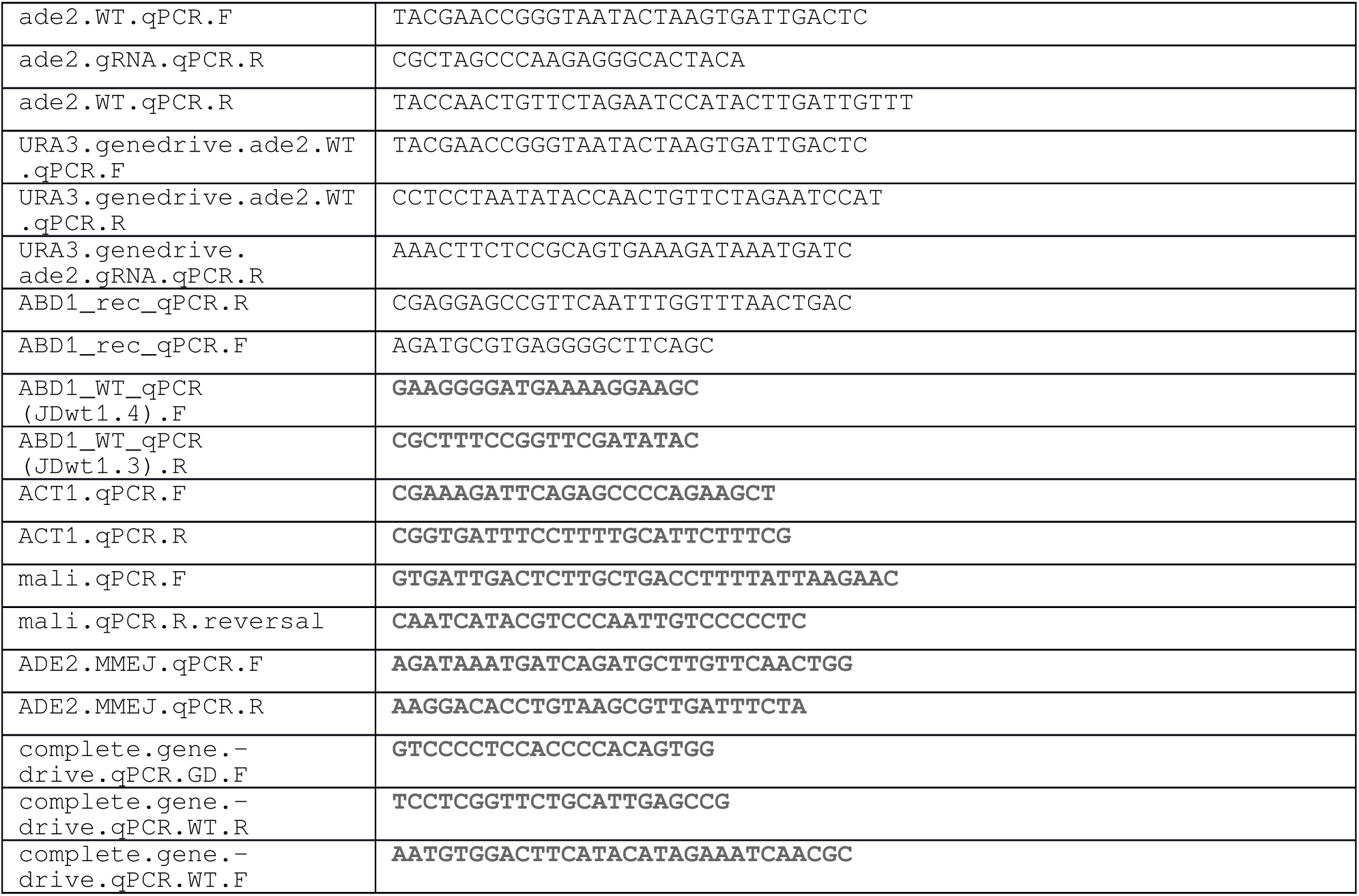

## Acknowledgements

We are very grateful to Steve Doris, Dan Spatt and Fred Winston for their incredible patience, generosity, and expertise in tetrad dissection. We also thank Fred Winston for providing us with SK1 strains and members of the Church laboratory for insightful discussions.

This work was supported by grants from the DOE (DE-FG02-02ER63445 to G.M.C.), NSF (SynBERC SA5283-11210 and MCB-1330914 to G.M.C.), NCI (5T32CA009216-34 to A.C.), NIDDK (1K99DK102669-01 to K.M.E.), and the Wyss Institute for Biologically Inspired Engineering (Technology Development Fellowship to K.M.E.).

## Author contributions

S.L.D. initiated the study; J.E.D., A.C., S.L.D., and K.M.E. designed the experiments; J.E.D. performed the experiments with assistance from A.C.; J.E.D., A.C., S.L.D., and K.M.E. analyzed the data; and K.M.E. wrote the paper with A.C. and contributing input from J.E.D., S.L.D., and G.M.C.

**Fig. 5-S1.**
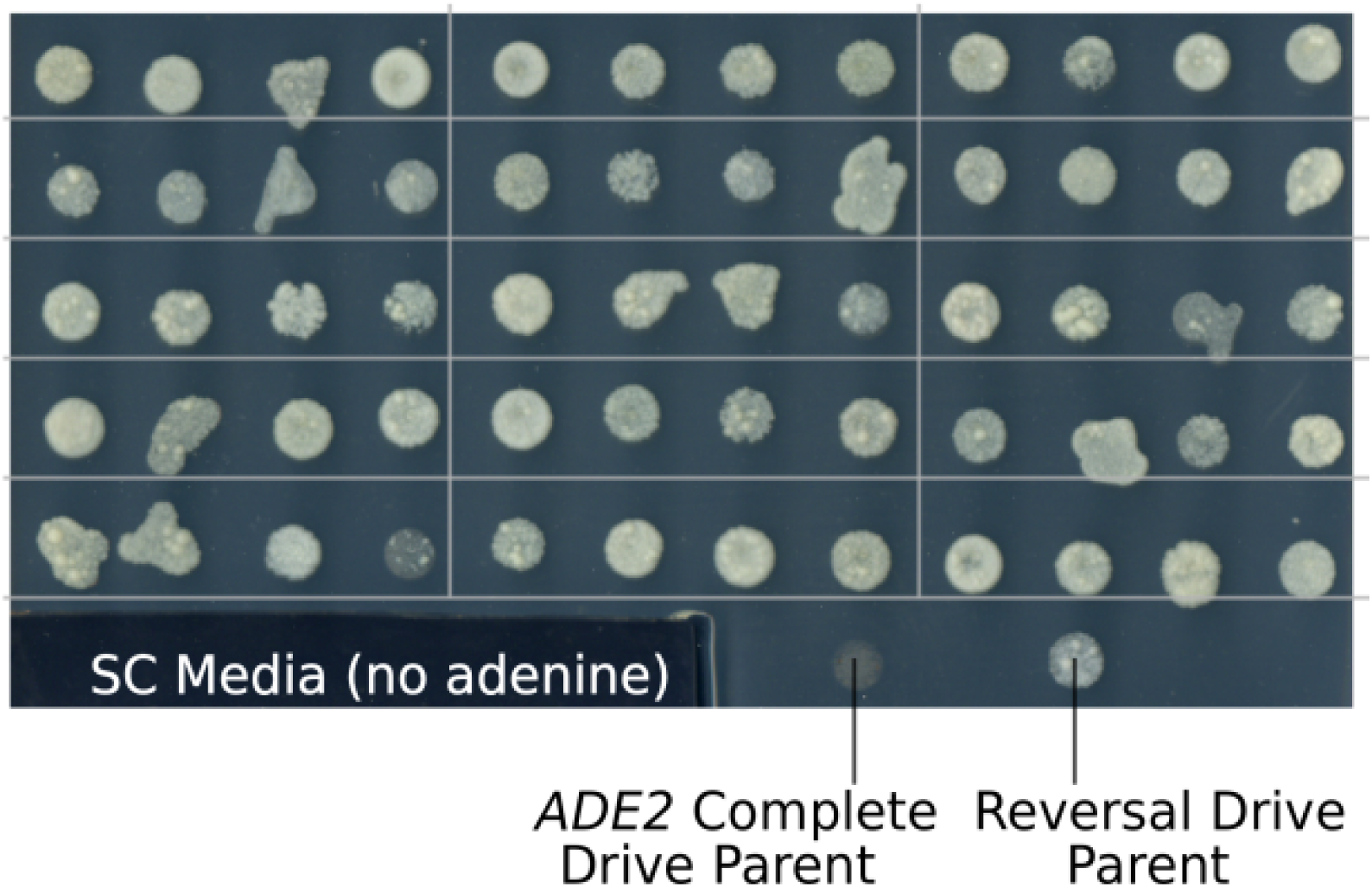
Reversal of drive-induced *ADE2* loss by a reversal drive. Haploid yeast containing a complete autonomous *ADE2*-disrupting gene drive were mated with haploids containing a reversal drive that restores *ADE2* function. 15 diploid offspring were sporulated, dissected, and plated on adenine-limited plates. The resulting cream-colored colonies indicate that an intact *ADE2* gene is present in all progeny, indicative of the reversal drive successfullycutting and replacing the *ADE2*-disrupting gene drive. (Return to Fig. 5)

